# In Silico Tool for Predicting and Scanning Rheumatoid Arthritis-Inducing Peptides in an Antigen

**DOI:** 10.1101/2025.03.19.644081

**Authors:** Ritu Tomer, Shipra Jain, Pushpendra Singh Gahlot, Nisha Bajiya, Gajendra P. S. Raghava

## Abstract

Rheumatoid arthritis (RA) is an autoimmune disorder in which the immune system mounts an abnormal response to self-antigens, leading to chronic inflammation and joint damage. Therefore, identifying antigenic regions in a protein that trigger RA is crucial for developing protein-based therapeutics. In this study, we developed models for predicting RA-inducing peptides using a dataset comprising of 291 experimentally confirmed RA-inducing peptides and 165 RA non-inducing peptides. Our initial analysis revealed that certain residues, such as glycine, proline, and tyrosine, are significantly enriched in RA-inducing peptides. While alignment-based techniques like BLAST and MERCI offered high precision, they suffered from limited coverage. We developed machine/deep learning based prediction and obtained highest performance (AUC = 0.75) using XGboost on an independent dataset. We also developed prediction methods using large language models and achieved highest performance (AUC 0.72) using ProtBERT. Our ensemble model achieved highest performance (AUC = 0.80 & MCC = 0.45) on an independent dataset that combine XGBoost and MERCI-derived motifs. All models were rigorously evaluated on an independent dataset not used during training or testing of models. This study will be valuable for assessing the risk of proteins used in probiotics, genetically modified foods, and protein-based therapeutics. Our most effective approach has been implemented in RAIpred, a web server and standalone software tool for predicting and scanning RA-inducing peptides. (https://webs.iiitd.edu.in/raghava/raipred/).

**Highlights:**

- Rheumatoid arthritis (RA), an incurable chronic joint disorder with diverse systemic complications.
- An attempt to identify antigenic regions in a protein which trigger this severe disease.
- Utilizing sequence composition based features for developing models.
- Implementation of ML, DL and LLM based models for prediction of RA-inducing peptides.
- Development of webserver, standalone, pypi and GitHub package for users.

## Introduction

Rheumatoid arthritis (RA) is an incurable chronic autoimmune joint disorder which exhibits significant clinical heterogeneity (Ding et al., 2023; Wasserman, 2011; J. Zhao et al., 2021). RA is characterized by abnormal inflammation within the joint’s synovial tissue, which progressively damages both cartilage and bone (Gibofsky, 2014; Smolen et al., 2016). The global prevalence of RA, affects approximately 1% of the world population, translating to millions of individuals (Huang et al., 2021; Ngian, 2010). In literature several studies have reported, that RA contributes to a diverse range of systemic complications like, cardiovascular disease (Gibofsky, 2014; D. Wu et al., 2022). The pathogenesis of RA is still unknown, but as reported it may arise from interactions among genetic predisposition and environmental factors. Several immune cells secrete immune regulatory molecules, which causes inflammation and joint damage, characteristic of autoimmune diseases (More et al., 2024). Previously, in silico methods has been developed to predict binders of HLA-DRB1*04:01 as it plays critical role in rheumatoid arthritis (Bhasin & Raghava, 2004; Patiyal et al., 2024).

In response to genetic and environmental triggers, autoreactive CD4+ T cells become activated, which presents antigens to B cells, producing autoantibodies such as rheumatoid factor and anti-citrullinated protein antibodies (C.-Y. Wu et al., 2021). This autoimmune response is further fuelled by pro-inflammatory cytokines, like tumor necrosis factor-alpha (TNF-α) and interleukin-6 (IL-6), which promote elevated immune cell response and inflammation in the synovium (McInnes & Schett, 2017). Additionally, macrophages and synovial fibroblasts releases the chemokines that perpetuates the inflammatory response (Hammaker et al., 2019). The dysregulation of the Janus kinase (JAK)/signal transducer and activator of transcription (STAT) signalling pathway is crucial, as it mediates the signalling of several key cytokine receptors involved in RA (Ding et al., 2023; Simon et al., 2021). Together, these pathways add in a persistent autoimmune response, leading to chronic joint inflammation and subsequent tissue destruction (Radu & Bungau, 2021).

In literature, traditional therapy for rheumatoid arthritis (RA) primarily focuses on disease-modifying anti-rheumatic drugs (DMARDs) (Radu & Bungau, 2021). These drugs have been reported to reduce the pro-inflammatory cytokine production, thereby decreasing the underlying inflammation process in synovium, which slows down the disease progression (Demoruelle & Deane, 2012). Notable progress has been observed in the DMARDs that target inflammation to prevent joint damage. Methotrexate is considered as first-line therapy response due to its efficacy and safety reported in literature (Bedoui et al., 2019; Cronstein, 1997; Z. Zhao et al., 2022). In addition to this, hydroxychloroquine and sulfasalazine are also widely considered conventional DMARDs, that can be administered alone or in combination with methotrexate (Conley et al., 2023; Moreira et al., 2022). Apart from DMARDs glucocorticoids, non-steroidal anti-inflammatory drugs (NSAIDs), and inflammatory cytokine inhibitors (ICI), are also used in the prevention of RA disease progression and management (J. Zhao et al., 2021). These traditionally used drugs have their own limitations that include inadequate response, intolerance, higher costs and number of side effects (Angelini et al., 2020).

In protein therapeutics era, one of the key challenge is identifying the antigens or antigenic regions that activate T-helper cells implicated in rheumatoid arthritis (Ansari et al., 2010). In the past, numerous computational methods have been developed to predict T-helper epitopes responsible for inducing cytokines such as interferon-gamma, TNF-alpha, IL-4, and IL-5 (Dhall et al., 2023, 2024; Dhanda et al., 2013; Naorem et al., 2023). These cytokines may play a crucial role in the development of autoimmune diseases like rheumatoid arthritis. Till date, there is no in-silico computational tool available in market, which aids in predicting T-helper cell-inducing peptides or epitopes that trigger rheumatoid arthritis. In this study, we extracted only the peptides that activate T-cells responsible for inducing rheumatoid arthritis. We extracted experimentally validated 291 RA associated and 165 non-associated peptides for IEDB. In order to create a robust model, we have implemented both alignment based like BLAST, MOTIF and alignment free approaches such as machine learning algorithms, deep learning, large language models. In addition, we developed an ensemble model that combines our best machine learning models with a motif-based approach to achieve higher performance. Finally, we developed a web server and standalone software RAIpred for predicting, designing and scanning RA inducing peptides.

## Materials & Methods

### 1. Dataset Preparation & Preprocessing

We gathered the experimentally validated data from the immune epitope database (IEDB) for our study and perform few preprocessing steps to further improve the quality of data used in the given study (Vita et al., 2019). Firstly, we extracted a total of 344 unique Rheumatoid arthritis (RA) inducing peptides as a positive dataset and 176 unique Rheumatoid arthritis (RA) non-inducing peptides (which are not available in positive) as negative dataset from IEDB. Here, we have observed that RA-associated peptides are HLA-class I and HLA-class II binders (please refer figure 2). The HLA-class-I contains only 46 peptides while HLA-class II have 298 peptides. Due to less number of peptides in HLA-class I, we have selected HLA-class II peptides only.

**Figure 1:**
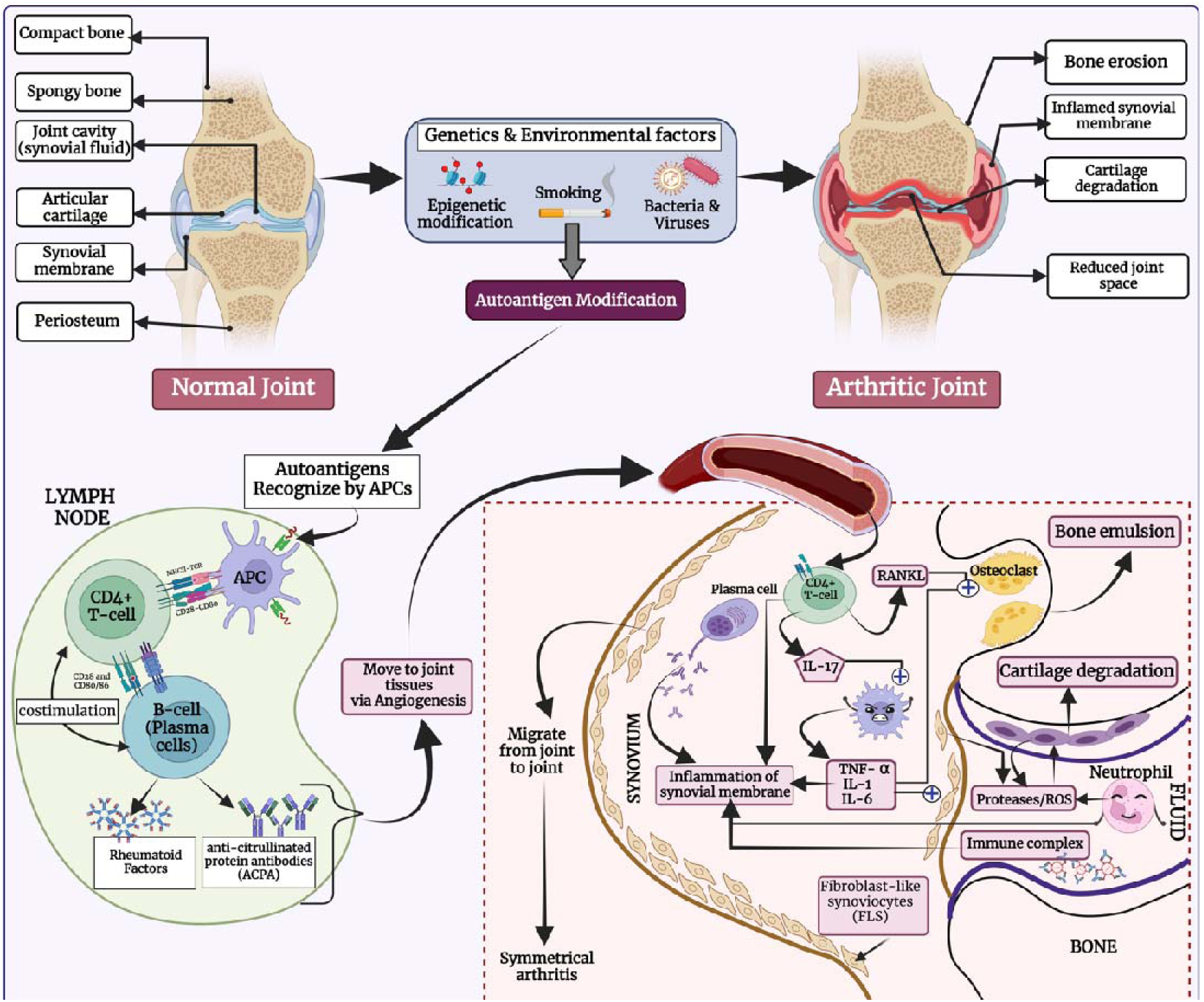
The figure depicts the etiology of rheumatoid arthritis.

**Figure 2:**
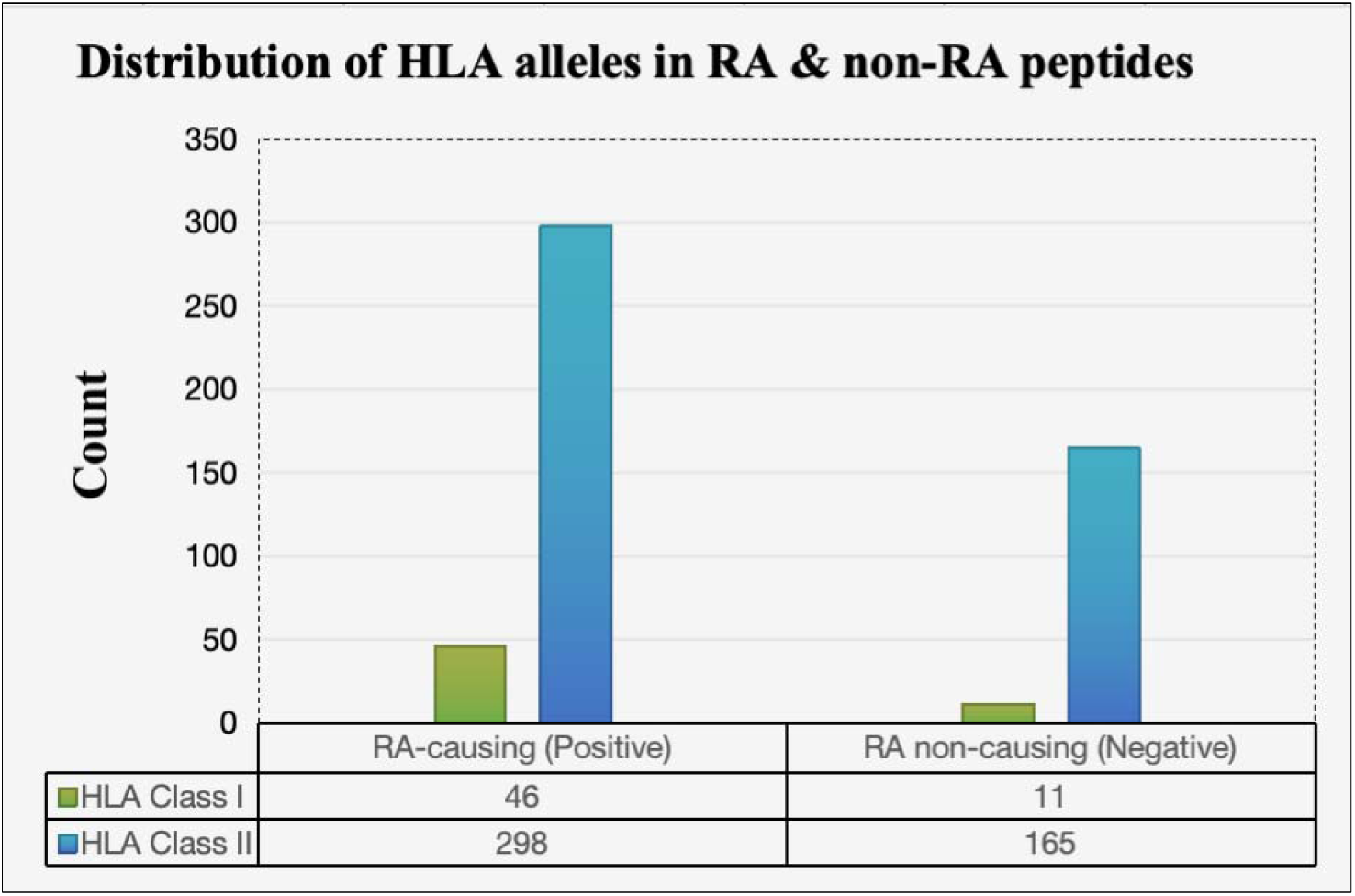
The figure shows the percentage of peptides binds to specific class/allele of MHC given in RA-inducers and RA non-inducers dataset.

**Figure 3:**
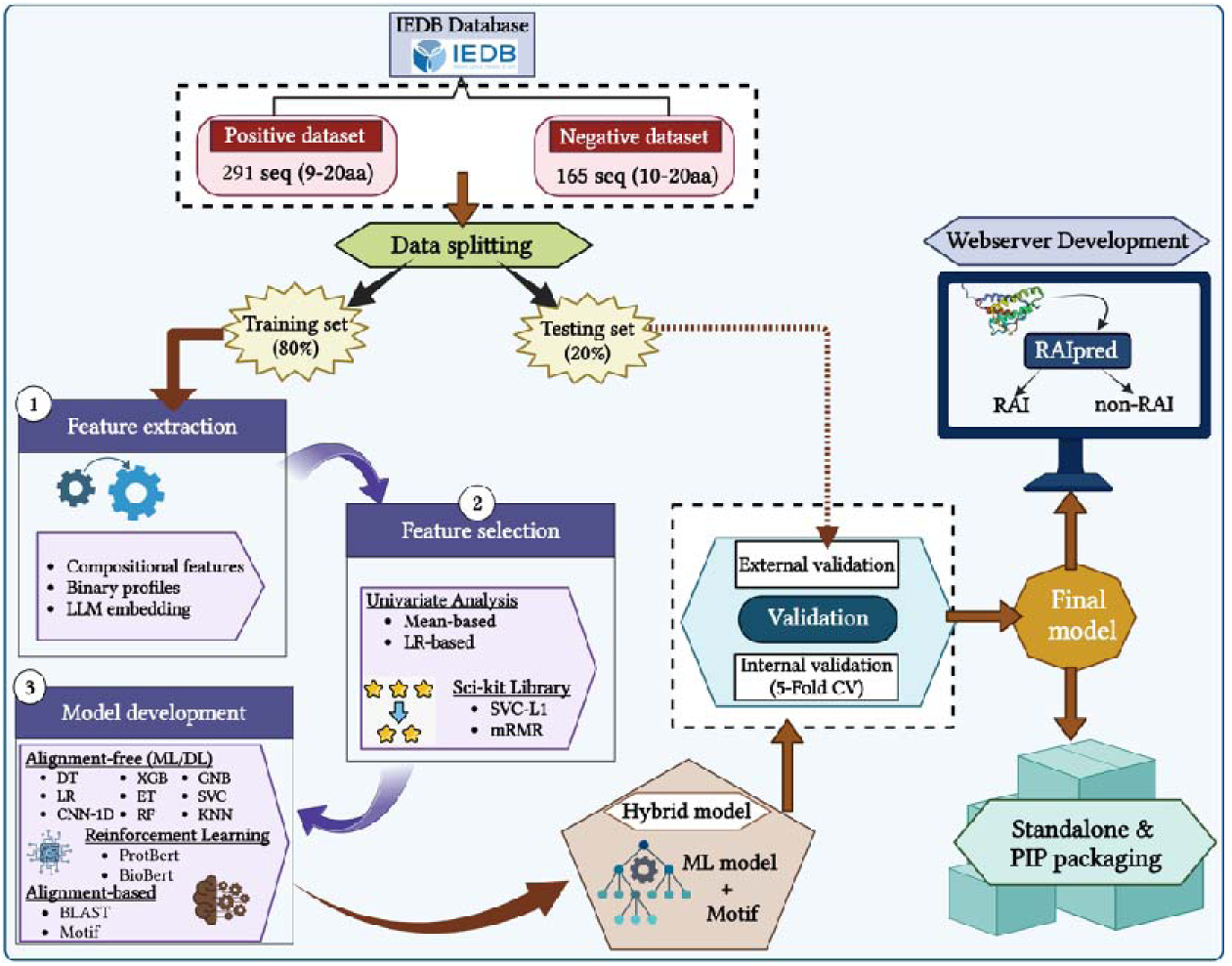
The figure depicts the complete operational flow of RAIpred.

Secondly, we perform few preprocessing steps such as removal of duplicate sequences from negative dataset which are common in both (Positive as well as negative) datasets. Then we filtered out sequences whose length occurrences are only 1 or few, we kept sequences more than 9 or less than 20 length of amino acid. After pre-processing of our data, we left with 291 sequences in positive dataset and 165 sequences in negative dataset.

### 2. Feature Generation

In present study, we have generated relevant features for both RA inducing as well as non-inducing peptides. We have calculated the different composition based features using Pfeature software (Pande et al., 2023). The Pfeature software calculates around 9189 features using amino-acid sequences. We have extracted features such as AAC, DPC, DDR, BTC, PCP, ATC, RRI, CeTD, PAAC, APAAC and many more. In addition to composition based features, we extend the features extraction to binary profiles as well. With the intent of capturing maximum information for the amino acid peptide sequences, we deployed the Amino Acid Binary Profile (AABP) module of Pfeature software (Pande et al., 2023). The generated features aids in implementing ML-based prediction algorithms. We have also included Embeddings of Large Language Models trained on ProtBERT developed by Rostlab (Ahmed Elnaggar, Michael Heinzinger, Christian Dallago, Ghalia Rihawi, Yu Wang, Llion Jones, Tom Gibbs, Tamas Feher, Christoph Angerer, Martin Steinegger, Debsindhu Bhowmik, 2020). Each and every feature have their own importance.

### 3. Preliminary Analysis

#### 3.1. Positional Analysis

We created a two sample logo implementing the Two Sample Logo software to comprehend the preference of amino acid residues at a certain location (Vacic et al., 2006). A fixed length input sequence vector is required for this step. We identified 9-mers from the N-terminal and 9-mers from the C-terminal of each peptide as the minimum length of peptides in our datasets is nine residues. Then, to produce a fixed length vector of 18 amino acids, we re-join both areas. For the purpose of creating one sample logo plot, 18-residue sequences were provided for each peptide in the positive and negative datasets.

#### 3.2. Compositional Analysis

To understand the deeper insights of amino acid composition difference between RA-inducing and non-inducing peptides, we perform compositional analysis on both positive and negative datasets. Here, we have utilized Pfeature software to compute amino acid compositions of both datasets (Pande et al., 2023). The Pfeature software calculates amino acid composition using formula;

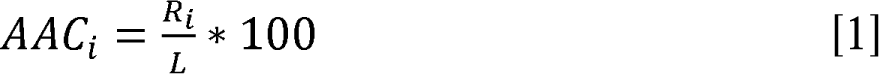

Where, AAC_i_ is amino-acid composition of residue type i, Ri is the number of residues in i, and L is the length of peptide sequence.

#### 3.3. Mean-based Univariate analysis

In mean based analysis, we have calculated the absolute mean difference of every feature in both positive and negative class after normalizing the data. In order to understand the relevance of feature generated, we computed the mean difference of each feature among both groups. Then we have applied the applied independent t-test and identified top 5 features with significant p-value among both groups.

#### 3.4. Logistic regression based analysis

We have also employed Logistic regression (LR) based single feature analysis. In this case, we have used LR as a statistical model to evaluate the relationship between each feature and the target label. We have computed AUC values for each feature to understand the relevance of each feature.

### 4. Alignment-based approach

#### 4.1. BLAST Search

We have utilised a similarity-search based well known method called BLAST for the annotation of peptide sequences (Altschul et al., 1990). In order to predict RA-inducing and RA non-inducing peptides, we used the peptide-peptide BLAST-based algorithm blastp-short (BLAST+ 2.2.28) in our study.

#### 4.2. Motif Search

Motifs are generally short amino acid patterns which might be responsible for a similar function. This analysis helps us to map signature patterns in inducing and non-inducing peptides. Motifs can act as targets to develop new drugs and therapies against RA. We deployed, Motif-EmeRging and with Classes-Identification (MERCI) tool (Vens et al., 2011) developed over Perl script to search for exclusive motifs in positive and negative sequences using default and user-defined parameters.

### 5. Alignment free approach

#### 5.1. Machine Learning Models

The Python software “scikit-learn” was used for classification. The classification algorithm consists of decision trees (DT), random forest (RF), multi-layer perceptron (MLP), eXtreme gradient boosting (XGBoost), support vector with the kernel as a radial basis (SVR), ExtraTreesClassifier (ET), Logistic Regression (LR), K-nearest Neighbor (KNN), and Guassian Naïve Baise (GNB). The grid search CV approach was used to tune hyperparameters, with “ROC” serving as the optimization metric.

#### 5.2. Deep Learning Models

In order to handle sequential data and extract hierarchical features along the sequence, we have used the 1D CNN model for the DL technique. The 1D CNN Model recognizes patterns and dependencies throughout the peptide sequences, making it especially useful for sequential data processing. By modifying hyperparameters, the model was individually adjusted on the datasets to maximize its performance for the classification task.

### 6. Feature Selection

As the literature suggests that not all descriptors are effective for developing machine learning models, two techniques were employed to select the best features: minimum Redundancy - Maximum Relevance (mRMR) and SVC-L1. Feature selection was performed on all combined composition based features. SVC-L1 based method able to select 34 features while we retrieve 50 features using mRMR based technique.

In addition to this, we have selected top relevant features from mean based univariate analysis and logistic regression based analysis. After computing the mean difference among both groups of each feature, we selected the 3782 out of 9189 features. Secondly, we applied independent t-test to identify significant features from the stretch of 3782 features using p-value <= 0.05. Finally, we have obtained 305 features with maximum absolute mean difference ranges between 0.001 to 0.14 with significant p-value. Upon which, we have deployed machine learning classifiers over top 10, 20, 50, 100, 150, 200, 250 and 305 features. Similarly, we have developed machine learning models on top 10, 20 and 50 features obtained from logistics based regression analysis.

### 7. Reinforcement learning

When it comes to text classification, LLM (large language model) excels because of their deep linguistic comprehension. LLMs can categorize data accurately even in situations where there is a lack of information because of their adaptability and remarkable capacity to extract important textual features. This study used LLMs to differentiate between RA inducing and RA non-inducing sequences. To fit the properties of the datasets, we used and fine-tuned the ProtBert model. After this fine-tuning, the model predicted whether each sequence belongs to the class of RA inducing and RA non-inducing.

### 8. Ensemble method

To develop a more robust model which can classify RA-inducing peptides from non-inducers, we apply two approaches to create an ensemble method: 1. Combination of BLAST based approach and ML-based method 2. Combination of motif based approach and ML-based method.

Firstly, BLAST based approach is used to identify disease causing peptides based on similarity hits and then machine learning approach is used for the prediction of those peptides which are not covered by the BLAST-based approach. Secondly, Motif-based approach is used to classify between disease causing peptides by identifying specific motifs and then machine learning approach is used for the prediction of those peptides which are not covered by the motif-based approach.

### 9. Model Evaluation

In present study, we have followed best machine learning practices, in order to avoid overfitting, we have implemented five-fold cross validation technique (Kumar et al., 2023; Tomer et al., 2023). The complete dataset was divided into 80:20 ratios, where 80% dataset used for the training and 20% used for the external validation. The model evaluation process was performed by employing standard parameters, here we estimated both threshold dependent and threshold independent parameters. In the case of threshold-dependent characteristics, we calculated sensitivity (Sens), specificity (Spec), accuracy (Acc), and Matthews correlation coefficient (MCC) using the equations (2–6). Furthermore, we evaluated the performance of models using a well-known and threshold-independent parameter Area Under the Receiver Operating Characteristic (AUROC) curve.

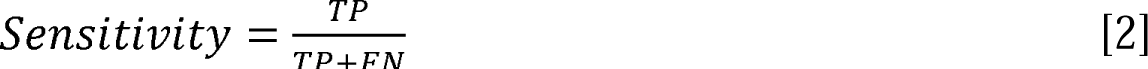

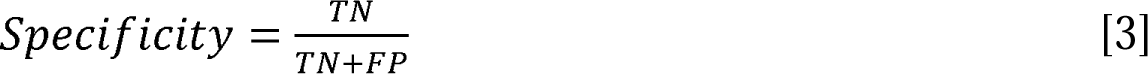

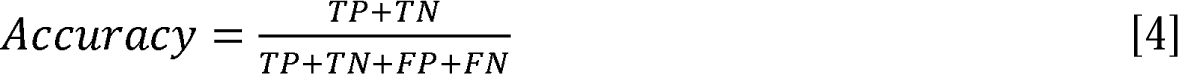

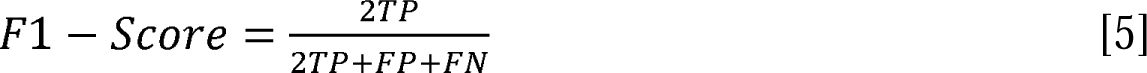

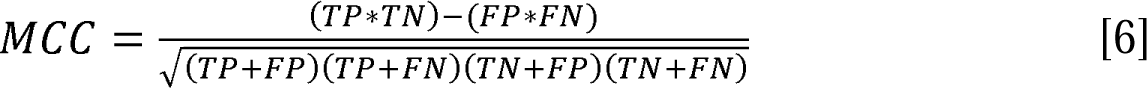

Where, FP is false positive, FN is false negative, TP is true positive and TN is true negative.

## Results

### 1. Preliminary Analysis

#### 1.1. Positional Analysis

To determine the most significant position of an amino acid residue in a peptide, we use Two Sample Logo to perform positional analysis. It is worth mentioning that the first nine regions relate to peptide N-terminus residues, while the last nine correspond to the peptide C-terminus. We discovered that Glycine (G), glutamine (Q), and phenylalanine (F) residues are located in the N-terminus regions of positive peptides, whereas Isoleucine (I), glycine (G), tyrosine (Y), and proline (P) are found in the C-terminus region. However, threonine (T) and isoleucine (I) are found at the N-terminus, and alanine (A), leucin (L), arginine (R), and histidine (H) are identified in the C-terminus area, whereas glutamic acid (E) highly prominent at every position in negative dataset.

#### 1.2. Compositional Analysis

Here, we computed amino acid composition for RA-inducing (positive) and RA non-inducing (negative) peptides. In figure 4, we can see that the glycine (G), proline (P) and tyrosine (Y) have the highest average composition in RA-inducing peptides with significant p-value as compared to the negative set. While the amino-acids alanine (A), aspartic acid (D), glutamic acid (E) and leucine (L) are significantly abundant in negative set.

**Figure 4:**
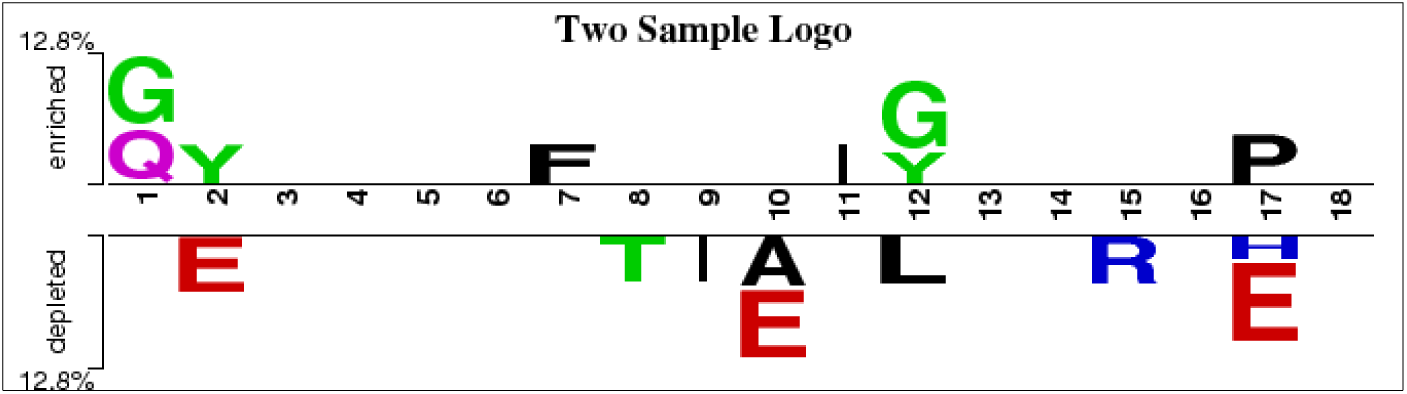
Positional analysis of RA inducing and non-inducing peptides generated using Two Sample Logo.

**Figure 5:**
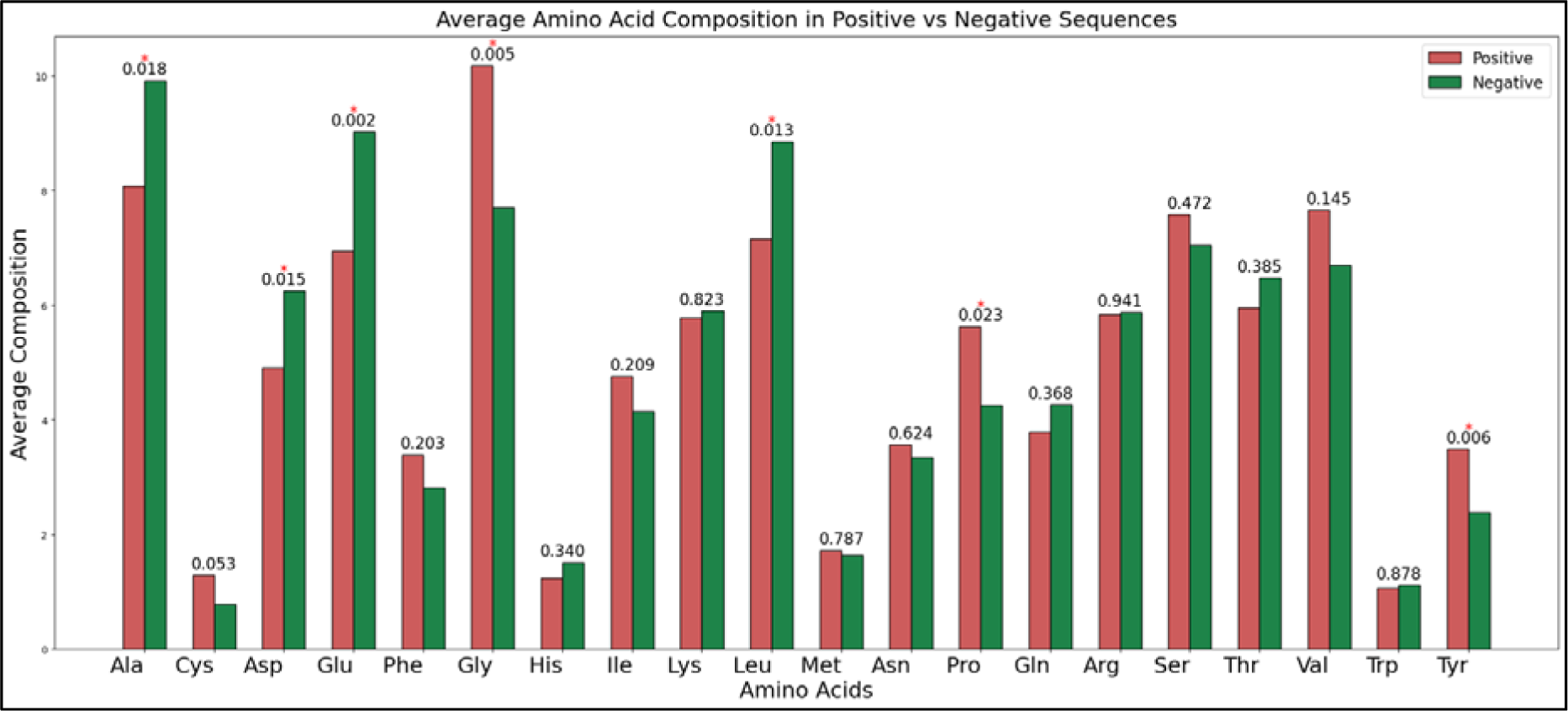
Amino acid compositional analysis among RA inducers and RA non-inducers.

#### 1.3. Mean-based univariate analysis

We have finally selected 305 features based on their higher mean difference between positive and negative dataset with significant p-value. As depicted in Table 2, the composition enhanced transition and distribution-based features such as CeTD_21_HB and CeTD_25_p_VW3 showed the highest mean difference of 0.084 & -0.140. Please refer **Supplementary Table S1** for the complete list and their rankings.

**Table 1:** The table shows list of features calculated using Pfeature tool.

| Type of Feature | Vector |
| --- | --- |
| AAC (Amino acid composition) | 20 |
| DPC (Dipeptide composition) | 400 |
| TPC (Tripeptide Composition) | 8000 |
| BTC (Bond composition) | 4 |
| DDR (Distance Distribution of Residues) | 20 |
| CTC (Conjoint triad calculation) | 343 |
| PRI (Repeat of Physico-chemical Properties) | 24 |
| CeTD (Composition enhanced Transition and Distribution) | 189 |
| APAAC (Amphiphilic pseudo amino acid composition) | 23 |
| ALLCOMP (All composition based features) | 9189 |
| AABP (Amino Acid Binary Profile) | 20*L |

**Table 2:** The table shows the top 5 features with maximum mean differences and a significant p-value.

| Feature | Pos_mean | Neg_mean | Diff | Feature | Pos_mean | Neg_mean | Diff |
| --- | --- | --- | --- | --- | --- | --- | --- |
| CeTD_21_HB | 0.466 | 0.382 | 0.084 | CeTD_25_p_VW3 | 0.306 | 0.445 | -0.140 |
| SER_Y | 0.275 | 0.196 | 0.079 | CeTD_100_p_VW3 | 0.320 | 0.453 | -0.133 |
| PRI_NE | 0.340 | 0.265 | 0.075 | SER_E | 0.382 | 0.514 | -0.132 |
| PRI_NE_pH | 0.340 | 0.265 | 0.075 | CeTD_75_p_VW3 | 0.340 | 0.471 | -0.131 |
| CeTD_11_SS | 0.373 | 0.299 | 0.074 | CeTD_50_p_VW3 | 0.330 | 0.458 | -0.128 |

#### 1.4. LR-based analysis

In order to identify the best features based on their individual performance, we have also applied logistic regression classifier on 9189 features, which were calculated using Pfeature tool. Here, we have observed that top AUROC features are from the bond composition (BTC) & Composition enhanced Transition and Distribution (CeTD) based feature categories. The features named as BTC_T, BTC_S and BTC_H achieved the maximum AUROC 0.69 & features CeTD_75_p_VW3, CeTD_100_p_VW3 and CeTD_50_p_VW3 achieved the AUROC of 0.68. The top 10 features with their performance are shown in **Table 3**, detailed results can be seen in **Supplementary Table S2.**

**Table 3:**
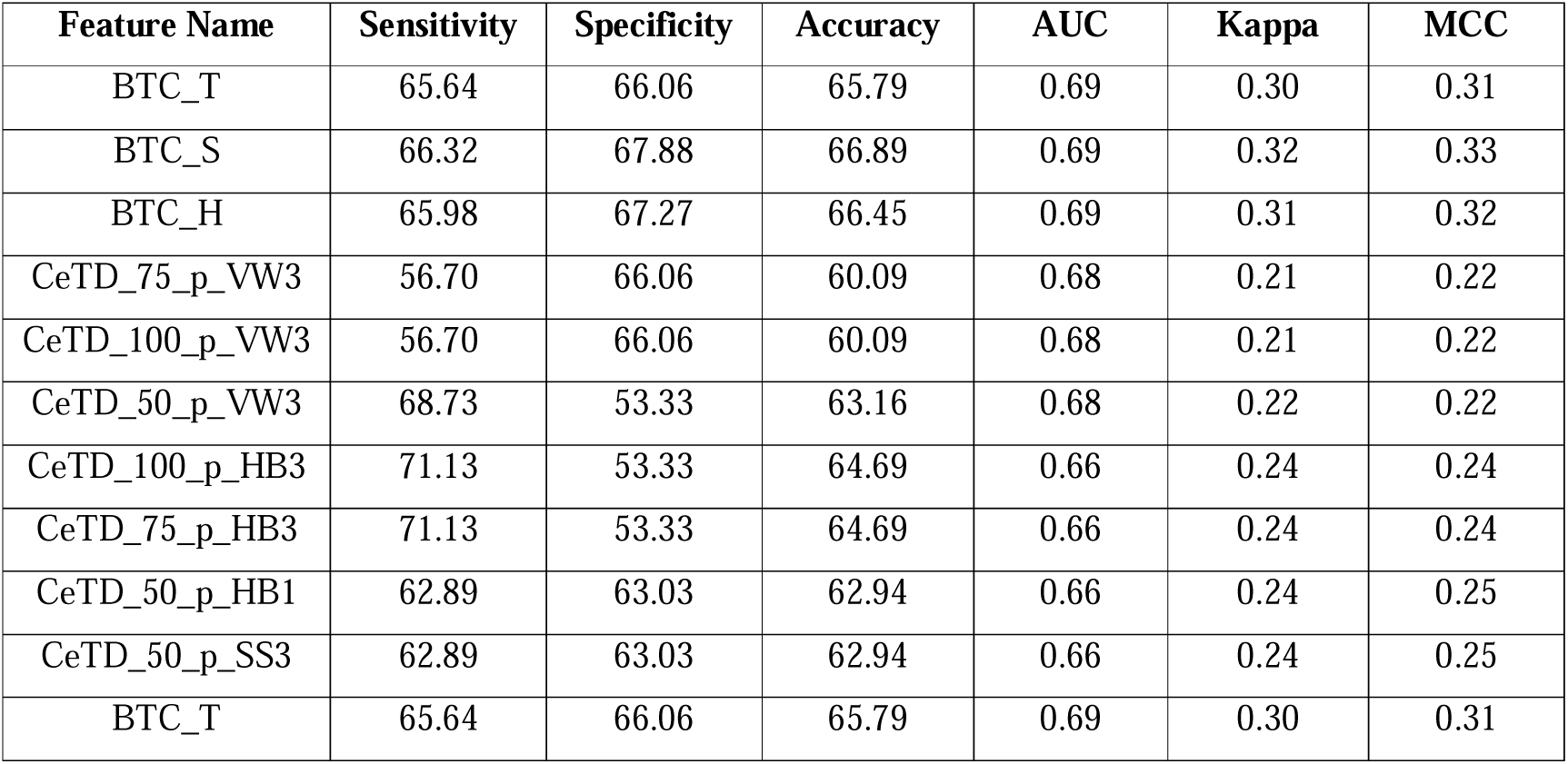
The table shows the performance of top 10 features using single feature analysis.

| Feature Name | Sensitivity | Specificity | Accuracy | AUC | Kappa | MCC |
| --- | --- | --- | --- | --- | --- | --- |
| BTC_T | 65.64 | 66.06 | 65.79 | 0.69 | 0.30 | 0.31 |
| BTC_S | 66.32 | 67.88 | 66.89 | 0.69 | 0.32 | 0.33 |
| BTC_H | 65.98 | 67.27 | 66.45 | 0.69 | 0.31 | 0.32 |
| CeTD_75_p_VW3 | 56.70 | 66.06 | 60.09 | 0.68 | 0.21 | 0.22 |
| CeTD_100_p_VW3 | 56.70 | 66.06 | 60.09 | 0.68 | 0.21 | 0.22 |
| CeTD_50_p_VW3 | 68.73 | 53.33 | 63.16 | 0.68 | 0.22 | 0.22 |
| CeTD_100_p_HB3 | 71.13 | 53.33 | 64.69 | 0.66 | 0.24 | 0.24 |
| CeTD_75_p_HB3 | 71.13 | 53.33 | 64.69 | 0.66 | 0.24 | 0.24 |
| CeTD_50_p_HB1 | 62.89 | 63.03 | 62.94 | 0.66 | 0.24 | 0.25 |
| CeTD_50_p_SS3 | 62.89 | 63.03 | 62.94 | 0.66 | 0.24 | 0.25 |
| BTC_T | 65.64 | 66.06 | 65.79 | 0.69 | 0.30 | 0.31 |

### 2. Alignment based approach

We used alignment-free methods (machine learning techniques) as well as alignment-based methods (Motif & BLAST), as shown in the previous sections. Every strategy has pros and cons of its own. Alignment-based techniques have a low sensitivity but a high specificity. The presence of motif or sequence similarity is essential to their performance. However, alignment-free approaches or models based on machine learning are more universal and performance is independent of similarity. We developed ensemble or hybrid approaches using BLAST & Motif to leverage the advantages of both alignment-free and alignment-based models.

#### 2.1. BLAST

In BLAST based approach Firstly, we have prepared a BLAST formatted database using training data. Then, we search query sequences (sequences in the test set) against the training database, to find hit at different e-values ranging from 1e-5 to 1e+3. The query sequence was categorized as positive if the top hit was positive and as negative if the top hit was negative. The detailed hits achieved by BLAST on validation data is shown below in **Table 4**.

**Table 4:** The table showing the coverage of BLAST hits on validation dataset.

| BLAST + CeTD (Validation) |  |  |  |  |  |
| --- | --- | --- | --- | --- | --- |
| E-value | Number of Hits | RA-inducers | RA non-inducers | Correct Pos | Correct Neg |
| **1.00E+10** | 92 | 46 | 46 | 10 | 23 |
| **1.00E+03** | 92 | 46 | 46 | 10 | 23 |
| **1.00E+02** | 87 | 44 | 43 | 9 | 20 |
| **1.00E+01** | 66 | 32 | 34 | 6 | 13 |
| **1.00E+00** | 53 | 28 | 25 | 6 | 12 |
| **1.00E-01** | **47** | **25** | **22** | **20** | **10** |
| **1.00E-02** | 39 | 21 | 18 | 4 | 10 |
| **1.00E-03** | 30 | 16 | 14 | 4 | 9 |
| **1.00E-04** | 20 | 12 | 8 | 4 | 6 |
| **1.00E-05** | 14 | 7 | 7 | 2 | 5 |
| **1.00E-08** | 2 | 1 | 1 | 0 | 1 |

Secondly, we have combined predicted labels developed using CeTD features with the BLAST score with the aim of enhancing the performance of our XGB models. We attained a maximum AUROC of 0.77 on the validation dataset at e-value 1.00E+01. As the e-value is too high so it could be the random chances of getting hits. The complete result table shown in **Supplementary Table S3.** In order to develop a more robust model, we further move-on to Motif-based approach.

#### 2.2. Motif

In Motif based approach, we have calculated K number of exclusive motifs using MERCI tool. The MERCI tool provides different parameters to generate specific motifs based on positive and negative sets. We have calculated exclusive motifs using None, KOOLMAN-ROHM and BETTS-RUSSELL classification methods and assign them +0.5 value if motif found in positive sequence and 0 if no match found. Finally, we combine predicted labels of best model with motif score. The best motifs with highest performance was obtained from BETTS-RUSSELL classification method. We have provided the detailed list of motifs with their occurrence in RA-inducing datasets in **Table 5**.

**Table 5:** The table shows the abundance of motifs in RA-inducing and RA non-inducing datasets.

| Motif | Coverage in RA-inducers |
| --- | --- |
| tiny polar hydrophobic G hydrophobic | 22 |
| hydrophobic L hydrophobic aliphatic small | 20 |
| hydrophobic polar hydrophobic tiny polar<br>hydrophobic hydrophobic polar | 19 |
| aliphatic aliphatic hydrophobic polar polar<br>aliphatic | 19 |
| small S hydrophobic G | 18 |
| hydrophobic polar A G hydrophobic | 17 |
| G small small G small | 17 |
| P hydrophobic polar polar hydrophobic | 17 |
| tiny hydrophobic S hydrophobic hydrophobic<br>hydrophobic | 16 |
| tiny S hydrophobic G | 15 |
| tiny S tiny tiny | 15 |

**Note:** polar : H,K,R,D,E,Y,W,T,C,S,N,Q; charged: D,E,R,H,K; negative: D,E; positive: R,H,K; small: A,G,C,S,P,N,D,T,V; tiny: A,G,C,S; hydrophobic: H,F,W,Y,I,L,V,M,K,T,A,G,C; aromatic: H,F,W,Y; aliphatic: I,L,V;

### 3. Alignment free Approach

#### 3.1. Machine Learning based analysis

We have applied multiple ML-based classifiers on composition based features and amino acid binary profile (AABP) based feature (Result given in **Supplementary Table S4**.), Our finding demonstrated that among different composition based features, the CeTD performed exceptionally. Using CeTD based features, we have achieved maximum accuracy and AUROC of 71% & 0.75 over training dataset and maximum accuracy and AUROC of 66.30% & 0.75 over validation dataset with balanced sensitivity and specificity by applying XGB classifier. The **Table 6**, below shows the performance of best model for all composition and binary profile based features over validation dataset.

**Table 6:** The table shows the performance of ML classifiers on validation dataset using composition and binary profile based features.

| Feature Name | ML Model | Sensitivity | Specificity | Accuracy | AUC | Kappa | MCC |
| --- | --- | --- | --- | --- | --- | --- | --- |
| CeTD | XGB | 61.02 | 75.76 | 66.30 | 0.75 | 0.33 | 0.35 |
| TPC | ET | 62.71 | 69.70 | 65.22 | 0.74 | 0.30 | 0.31 |
| ALLCOMP | LR | 57.63 | 75.76 | 64.13 | 0.73 | 0.30 | 0.32 |
| APAAC | ET | 61.02 | 63.64 | 61.96 | 0.72 | 0.23 | 0.24 |
| DPC | KNN | 61.02 | 69.70 | 64.13 | 0.71 | 0.28 | 0.30 |
| AAC | ET | 59.32 | 60.61 | 59.78 | 0.70 | 0.19 | 0.19 |
| BTC | GNB | 50.85 | 75.76 | 59.78 | 0.69 | 0.23 | 0.26 |
| DDR | ET | 52.54 | 75.76 | 60.87 | 0.67 | 0.25 | 0.28 |
| CTC | SVC | 62.71 | 66.67 | 64.13 | 0.65 | 0.27 | 0.28 |
| PRI | LR | 55.93 | 69.70 | 60.87 | 0.63 | 0.23 | 0.25 |
| AABP | KNN | 55.93 | 51.52 | 54.35 | 0.59 | 0.07 | 0.07 |

**Note:** XGB: extreme gradient boosting, ET: extra tree, LR: logistic regression, KNN: k-nearest neighbour, GNB: Guassian Naïve Baise, SVC: support vector classifier, AUC: area under curve, kappa: Cohen’s kappa coefficient, MCC: Mathew’s correlation coefficient.

#### 3.2. Deep Learning based analysis

We have applied 1DCNN on different composition-based features as well as on amino acid binary profile-based features. Here, 1DCNN, performs well on tri-peptide composition based features by achieving AUROC of 0.69 on validation dataset. The detailed results of 1DCNN model on different features are given in **Supplementary Table S5.** The **Table 7** of all feature performance is given below.

**Table 7:** The table shows the performance of 1DCNN model over different type of features on validation data.

| Feature Name | Sensitivity | Specificity | Accuracy | AUC | Kappa | MCC |
| --- | --- | --- | --- | --- | --- | --- |
| AAC | 88.14 | 21.21 | 64.13 | 0.60 | 0.11 | 0.13 |
| DPC | 50.85 | 63.64 | 55.44 | 0.67 | 0.13 | 0.14 |
| **TPC** | **55.93** | **72.73** | **61.96** | **0.69** | **0.26** | **0.28** |
| BTC | 0.00 | 100.00 | 35.87 | 0.38 | 0.00 | 0.00 |
| DDR | 55.93 | 45.46 | 52.17 | 0.54 | 0.01 | 0.01 |
| CTC | 52.54 | 66.67 | 57.61 | 0.60 | 0.17 | 0.19 |
| PRI | 45.76 | 69.70 | 54.35 | 0.64 | 0.14 | 0.15 |
| CeTD | 71.19 | 48.49 | 63.04 | 0.68 | 0.20 | 0.20 |
| APAAC | 59.32 | 51.52 | 56.52 | 0.65 | 0.10 | 0.11 |
| AAB | 55.93 | 54.55 | 55.44 | 0.60 | 0.10 | 0.10 |
| **ALLCOMP** | **57.63** | **69.70** | **61.96** | **0.72** | **0.25** | **0.26** |

**Note:** AUC: area under curve, kappa: Cohen's kappa coefficient, MCC: Mathew's correlation coefficient.

### 3.3. Feature selection techniques

In order to select most relevant features, we have applied two feature selection techniques – SVC-L1 and mRMR. Using SVC-L1 technique, we were able to select 34 different composition based features. As depicted in **Table 8**, we achieved maximum AUROC of 0.72 on validation dataset using support vector classifier (SVC).

**Table 8:** The table shows the performance of feature selected using SVC-L1 technique over validation dataset.

| Model | Sensitivity | Specificity | Accuracy | AUC | Kappa | MCC |
| --- | --- | --- | --- | --- | --- | --- |
| **DT** | 62.71 | 48.49 | 57.61 | 0.58 | 0.11 | 0.11 |
| **RF** | 59.32 | 72.73 | 64.13 | 0.70 | 0.29 | 0.31 |
| **LR** | 59.32 | 66.67 | 61.96 | 0.71 | 0.24 | 0.25 |
| **XGB** | 66.10 | 69.70 | 67.39 | 0.69 | 0.34 | 0.34 |
| **KNN** | 61.02 | 69.70 | 64.13 | 0.71 | 0.28 | 0.30 |
| **GNB** | 62.71 | 66.67 | 64.13 | 0.68 | 0.27 | 0.28 |
| **ET** | 69.49 | 54.55 | 64.13 | 0.69 | 0.24 | 0.24 |
| **SVC** | **62.71** | **75.76** | **67.39** | **0.72** | **0.35** | **0.37** |
| **MLP** | 62.71 | 66.67 | 64.13 | 0.71 | 0.27 | 0.28 |

**Note:** DT: decision tree, RF: random forest, LR: logistic regression, XGB: extreme gradient boosting, KNN: k nearest neighbour, GNB: Guassian Naïve Baise, ET: extra tree, SVC: support vector classifier, MLP: multilayer perceptron, AUC: area under curve, kappa: Cohen's kappa coefficient, MCC: Mathew's correlation coefficient.

In present study, we have also applied mRMR based feature selection technique, which were calculated 50 different composition based features and achieved a maximum AUROC of 0.73 on validation dataset using random forest (RF) classifier. The detailed results are depicted in **Table 9**.

**Table 9:** The table shows the performance of feature selected using mRMR based technique over validation dataset.

| Model | Sensitivity | Specificity | Accuracy | AUC | Kappa | MCC |
| --- | --- | --- | --- | --- | --- | --- |
| DT | 55.93 | 66.67 | 59.78 | 0.65 | 0.21 | 0.22 |
| RF | **59.32** | **69.70** | **63.04** | **0.73** | **0.27** | **0.28** |
| LR | 66.10 | 75.76 | 69.57 | 0.72 | 0.39 | 0.40 |
| XGB | 55.93 | 75.76 | 63.04 | 0.71 | 0.28 | 0.31 |
| KN | 57.63 | 72.73 | 63.04 | 0.69 | 0.27 | 0.29 |
| GNB | 79.66 | 39.39 | 65.22 | 0.60 | 0.20 | 0.21 |
| ET | 52.54 | 72.73 | 59.78 | 0.74 | 0.22 | 0.24 |
| SVC | 66.10 | 69.70 | 67.39 | 0.71 | 0.34 | 0.34 |
| MLP | 64.41 | 66.67 | 65.22 | 0.67 | 0.29 | 0.30 |

**Note:** DT: decision tree, RF: random forest, LR: logistic regression, XGB: extreme gradient boosting, KNN: k nearest neighbour, GNB: Guassian Naïve Baise, ET: extra tree, SVC: support vector classifier, MLP: multilayer perceptron, AUC: area under curve, kappa: Cohen's kappa coefficient, MCC: Mathew's correlation coefficient.

In addition to this, we have also implemented mean based univariate analysis and logistics regression based feature selection techniques. We have applied machine learning techniques on 305 features as well as on top 200, 150, 100, 50, 20, 10 set of features. Using this, we achieved highest AUC as 0.73 for top 100, top 150 and top 200 features. The results of all the selected features over validation dataset are shown below in **Table 10**, for detailed results please refer **Supplementary Table S6.** Also, the performance of various machine learning algorithms on top 10, 20 and 50 features selected using logistics based regression analysis to discriminate RA-inducers from non RA-inducers, can be referred in **Supplementary Table S7.**

**Table 10:** The table shows the best performing models over validation dataset on different set of features selected using Mean-based univariate analysis.

| Total Feature | ML Model | Sensitivity | Specificity | Accuracy | AUC | Kappa | MCC |
| --- | --- | --- | --- | --- | --- | --- | --- |
| Top 305 | RF | 57.63 | 66.67 | 60.87 | 0.72 | 0.22 | 0.23 |
| Top 200 | RF | 66.10 | 69.7 | 67.39 | 0.73 | 0.34 | 0.34 |
| **Top 150** | **RF** | 59.32 | 72.73 | 64.13 | 0.73 | 0.29 | 0.31 |
| **Top 100** | **ET** | 67.80 | 66.67 | 67.39 | 0.73 | 0.33 | 0.33 |
| **Top 50** | **ET** | 61.02 | 66.67 | 63.04 | 0.70 | 0.26 | 0.27 |
| **Top 20** | **SVC** | 50.85 | 75.76 | 59.78 | 0.70 | 0.23 | 0.26 |
| **Top 10** | **SVC** | 59.32 | 69.70 | 63.04 | 0.69 | 0.27 | 0.28 |

**Note:** RF: random forest, ET: extra tree, SVC: support vector classifier

### 4. Reinforcement learning based analysis

We have used protBERT and BioBERT, two pre-trained large language models, for this study. The number of epochs was changed to fine-tune the models for specific tasks. Among the epochs tested, epoch 10 on protBERT & 3 on BioBERT provided the best results, with a maximum AUC of 0.71 and 0.67 on finetune models. The **Table 11** shows the finetune model result over validation dataset.

**Table 11:** The table shows the performance of finetune LLM models over validation dataset.

| Model Name | Sensitivity | Specificity | Accuracy | AUC | MCC |
| --- | --- | --- | --- | --- | --- |
| **ProtBERT** | 0.81 | 0.58 | 0.73 | 0.71 | 0.40 |
| **BioBERT** | 0.69 | 0.58 | 0.65 | 0.67 | 0.26 |

Furthermore, since embeddings generated by the fine-tuned model can serve as valuable features, we have extracted embeddings using the above finetune model and applied various ML algorithms. The finetune model were not performed well on this data. The results for LLM models are given in **Supplementary Table S8.**

### 7. Ensemble Model

An ensemble model is a combined model developed by merging two different approaches to enhance the model quality and robustness. Here, we have combined motif-based approach with best performing ML model using CeTD based features. We have observed that the BETTS-RUSSELL classification method of MERCI with fp-2, fn-0, g-0, k-20 parameters identify the best motifs which covers maximum validation data. We have combined these motif scores with best performing ML-classifier. As depicted in **Table 12**, we achieved the highest AUROC of 0.80 with MCC of 0.45 on validation dataset. The exclusive motif of disease-inducing peptides would help easily identifying the specific regions in protein which leads to this disease. Finally. This ensemble model is also incorporated in the RAIpred webserver’s predict module for the providing the easy accessibility.

**Table 12:** The table shows the performance of hybrid model developed using exclusive positive motifs [with MERCI classification – None, KOOLMAN-ROHM and BETTS-RUSSELL].

| Classification Method | Requested Number of Motif | Sensitivity | Specificity | Accuracy | AUC | Kappa | MCC |
| --- | --- | --- | --- | --- | --- | --- | --- |
| None | K20 | 67.80 | 66.67 | 67.39 | 0.70 | 0.33 | 0.33 |
| KOOLMAN-ROHM | K20 | 71.19 | 66.67 | 69.57 | 0.75 | 0.36 | 0.37 |
| BETTS-RUSSELL | K20 | 71.19 | 75.76 | 72.83 | 0.80 | 0.44 | 0.45 |

**Note:** K: number of requested motifs, AUC: area under curve, kappa: Cohen's kappa coefficient, MCC: Mathew's correlation coefficient.

### 8. Webserver Designing

We developed RAIpred, which is accessible at https://webs.iiitd.edu.in/raghava/raipred/, to provide the scientific community a more user-friendly webserver. This platform employs our best-performing model to forecast peptides that cause RA. RAIpred provides “Prediction, Design, Protein Scan, and Motif Scan” modules. By efficiently differentiating between RA-inducing and RA-non-inducing proteins, the Prediction module enables users to submit protein sequences in FASTA format for prediction. Users can forecast which analogs will cause RA by using the Design module to produce all possible analogs of the input sequence. The Protein Scan module helps find the parts of a given protein sequence that cause RA. The “MERCI” program is used in the Motif Scan module to map the motifs in the query sequence that cause RA. The responsive framework used in the development of this platform allows it to be viewed on a variety of platforms, including computers, laptops, and smartphones. Furthermore, we have developed a standalone Python tool called “RAIpred” to help users identify areas of proteins or peptides that may cause sickness. The web server’s “download module” can be used to get this package at https://webs.iiitd.edu.in/raghava/raipred/download.html. Powered by HTML5, Java, CSS3, and PHP scripts, the webserver works with a number of devices, including desktop, tablet, mobile, and iMac.

## Discussion

Rheumatoid arthritis is caused by gradual loss of self-tolerance in genetically vulnerable individuals due to various environmental stressors. Both the genetic as well as environment factors are significantly responsible for the onset of the disease. The ability of peptides to selectively bind arthritogenic peptide sequences for presentation to auto-reactive T lymphocytes is provided by a common epitope in the peptide binding groove area of MHC class II molecules (Hill et al., 2003). These T-cells generate inflammatory responses by releasing excessive amounts of cytokines and by activating B-cells which are further responsible for the excessive production of autoantibodies and lead to destruction and bone erosion. As these arthritogenic peptides are responsible for inducing T-cell response, they can be targeted for therapeutic purposes by modifying their properties using peptide cyclization, chemical modification or other in-vitro approaches.

Few machine learning-based techniques have been developed in the past to combat rheumatoid arthritis. One of these techniques, which was developed in order to forecast how well biologic drugs will work in treating patients with RA and AS (ankylosing spondylitis), with an AUC of 0.64 based on validation data. Furthermore, it demonstrates that the most significant predictors of therapy responses were patient self-reporting scales, the Bath Ankylosing Spondylitis Functional Index (BASFI) in AS patients, and the patient global assessment of disease activity (PtGA) in RA patients (Lee et al., 2021). Another study uses machine learning (ML) to predict patient relapses based on blood test results and ultrasound examination data (Matsuo et al., 2022). Prasad et al. developed ATRPred, an ML-based technique that uses clinical and demographic characteristics to predict how well RA patients will respond to anti-TNF treatment (Prasad et al., 2022). One study uses genetic information from SNPs in non-HLA genes to predict RA (Dudek et al., 2024). To the best of our knowledge, no method has yet been developed to anticipate the T-helper cell-inducing peptides or epitopes that trigger rheumatoid arthritis.

In this study, we have made a systemic approach for the classification of RA-inducing peptides. We have extracted 291 RA-inducing peptides and 165 RA non-inducing peptides from IEDB. In the preceding sections, we observed few key insights from the comprehensive analysis among RA inducing and non-inducing peptides. Here, compositional analysis indicates that RA-inducing peptides have the highest average composition of glycine and proline as compared to non-inducing peptides, which might be responsible for peptide binding to MHC (Rauscher et al., 2006). In addition to this, positional analysis further marks the preferences of distinct amino acid preferences for N-terminal and C-terminal regions in positive and negative peptide datasets, emphasizing the differential roles of residues such as glycine (G), threonine (T), and alanine (A). In classification modelling, both alignment-based (BLAST and Motif) and alignment-free methods were implemented. BLAST based approach demonstrated slightly increased performance at higher e-value which depicts the random chances of getting hits. While motif based approach, gives highest correct hits on validation set. In the present study, among different composition-based features, we have observed that CeTD composition-based features outperformed all. We have reported maximum accuracy and AUROC over training data as 71% & 0.75 and 66.30% & 0.75 over validation dataset with balanced sensitivity and specificity by applying XGB classifier. This result underscores that CeTD features capture physicochemical peptide properties for our dataset, which is critical for accurate predictions. Finally, we have combined motif analysis and an ML-based methodology to develop an ensemble method. On the validation dataset, an ensemble-based method gets a maximum AUROC of 0.80 and an MCC of 0.45. The integration of compositional insights and machine learning algorithms enabled the development of a robust tool ‘RAIPred’ for RA-inducing peptide prediction. In order to server the scientific community an easy and user-friendly approach, we have also developed a web server named RAIpred (https://webs.iiitd.edu.in/raghava/raipred/), standalone package (https://webs.iiitd.edu.in/ raghava/raipred/download.html) and Github (https://github.com/raghava/raipred) for the prediction of RA-inducing peptides.

## Applications of RAIpred

- This tool enables researchers to access RA causing risk involved in newly discovered proteins/ peptides, before deploying them in therapeutics or GMOs.
- This tool aids in designing therapeutic peptides by assessing their physiochemical properties as well as screening them as RA inducers or non-inducers.
- Using Protein scan module users can map disease causing antigenic epitopes in the query protein sequence.
- Researchers may produce new peptides by using the Design Module of RAIpred to execute single residue modifications and anticipate whether or not they would accelerate the pathogenesis of RA.

## Conclusion

Nowadays, peptide-based therapeutics has become popular as they have high success rate in clinical trials due to their selectivity towards the target. Risk assessment of these proteins is essential for preventing them causing some severe side-effects or involve in disease development. There are several peptide-based drugs, approved by FDA for treatment of rheumatoid arthritis and other autoimmune disorders. This RA-inducing peptide prediction tool, with high accuracy and balanced sensitivity and specificity, making it a reliable resource for identifying RA-inducing peptides. This tool would aid in development of targeted therapeutic peptides treatment for RA. Additionally, it provides valuable insights into peptide functionality, aiding in the discovery of novel peptides with potential biological and pharmaceutical significance.

## Author Contributions

RT and GPSR collected and processed the datasets. RT and GPSR implemented the algorithms and developed the prediction models. RT, SJ and GPSR analysed the results. RT and PSG created the front-end user interface and RT created the back-end of the webserver. RT, SJ and GPSR penned the manuscript. NB prepared the figures for the manuscript. RT, SJ, PSG, NB and GPSR reviewed the manuscript. GPSR conceived and coordinated the project. All authors read and approved the final manuscript.

## Acknowledgments

Authors are thankful to the Department of Science and Technology (DST-INSPIRE), University Grants Commission (UGC), Department of Bio-Technology (DBT), and Council of Scientific & Industrial Research (CSIR) for fellowships and financial support, and the Department of Computational Biology, IIITD New Delhi for infrastructure and facilities. We would like to acknowledge that Figures were created using BioRender.com.

## Data Availability Statement

All the datasets used in this study are available at the “RAIpred” webserver, https://webs.iiitd.edu.in/raghava/raipred/dd.html.

## Abbreviations

RA: Rheumatoid arthritis
IEDB: Immune epitope database
HLA: Human leukocyte antigen
XGBoost: extreme gradient boosting classifier
ProtBERT: Protein BERT
DMARDs: Disease-modifying anti-rheumatic drugs
NSAIDs: Non-steroidal anti-inflammatory drugs
GMO: Genetically modified organisms

